# Bioinformatics Approach on Hypothetical Protein from *Clostridioides difficile*: Structural, Functional and Molecular Docking Analysis

**DOI:** 10.1101/2025.03.20.644477

**Authors:** David Jones

## Abstract

*Clostridioides difficile* (*C. difficile*) has become a globally important pathogen as epidemic strains spread through hospitals in many countries. Despite the advancements in infection therapy, there is a need for more efficacious medicines against *C. difficile* that simultaneously minimize injury to the resident gut microbiota. The study aims to investigate various aspects of the protein, including its physicochemical properties, subcellular location, functional elucidation, protein-protein interactions, structure prediction, validation, determination of active sites for potential ligands, and MD study. The protein is partially basic and hydrophobic, according to the physicochemical properties analysis. The protein has activities in the inner membrane and the cytoplasm with two transmembrane helices. Furthermore, the protein is involved in secondary transporters in the MFS system, resulting in the movement of various substances through cytoplasmic or internal membranes. We targeted the active sites of the protein as potential binding sites for ligand molecules to discover novel therapeutic agents. The MD study documented the interaction of the selected ligands (PAβN and CCCP) with the protein. PAβN demonstrated the most suitable ligand compared to CCCP, as it required the lowest energy (– 6.7 kcal/mol) to interact with the protein. This functional protein can be targeted for further study on potential therapeutic candidates against the protein of *C. difficile*.

## 1. Introduction

*C. difficile*, an anaerobic-toxigenic bacteria, induces severe infectious colitis, resulting in considerable morbidity and mortality globally. Both augmented bacterial toxins and weakened host immune responses trigger symptomatic illness. *C. difficile* has been a recognized pathogen in Europe and North America for decades and is also emerging in Asia [1]. Individuals are at greater risk for *C. difficile* infection, including people in hospitals over the age of 65 who have recently undergone treatment with antibiotics [2]. Antibiotics reduce protective gut flora, while age along with medical comorbidities decreases the immunological response to *C. difficile*. Hospital environments and long-term care settings typically host most epidemics, with occasional documentation of outpatient acquisition [3]. Even in the absence of past antibiotic exposure, the advent of hypervirulent strains in Europe and North America has widened the effect of *C. difficile* to include younger individuals and a larger community-based population [4].

The capacity of *C. difficile* to provoke enteritis relies on two host factors: colonization resistance and the immune response (IR) to *C. difficile*. The fecal microbiome, with around 4,000 bacterial species, safeguards the large intestine against invasive illnesses. [5]. These microorganisms give colonization resistance to disease-causing organisms by competing for critical nutrition and adhesion sites on the gut wall. Antibiotics (ABs) disrupt the barrier microbiota and reduce colonization resistance, increasing the development of gut infections [6]. It seems that drugs have a big effect on the pathophysiology of *C. difficile* because they lower the numbers of Firmicutes phyla and Bacteroides [7]. The fecal microbiota of neonates and infants has a deficiency in colonization resistance. Thus, around 60-70% of healthy babies are asymptomatic carriers of *C. difficile* throughout the first year of life [8]. Furthermore, patients who developed a suitable antibody response during an initial *C. difficile* infection episode exhibited a reduced risk of recurrence. In contrast, Solomon et al. established that individual with diminished serum antitoxin-A-IgG levels had a markedly higher likelihood of mortality within the initial 30 days of illness [9]. The immune system of adults is usually weaker against *C. difficile* because they are older, malnourished, or have other health problems. These factors may also be linked to more severe infections [9, 10].

## 2. Materials and methods

Figure **1** show the overall methodology and bioinformatics tools used in this work.

**Figure 1.**
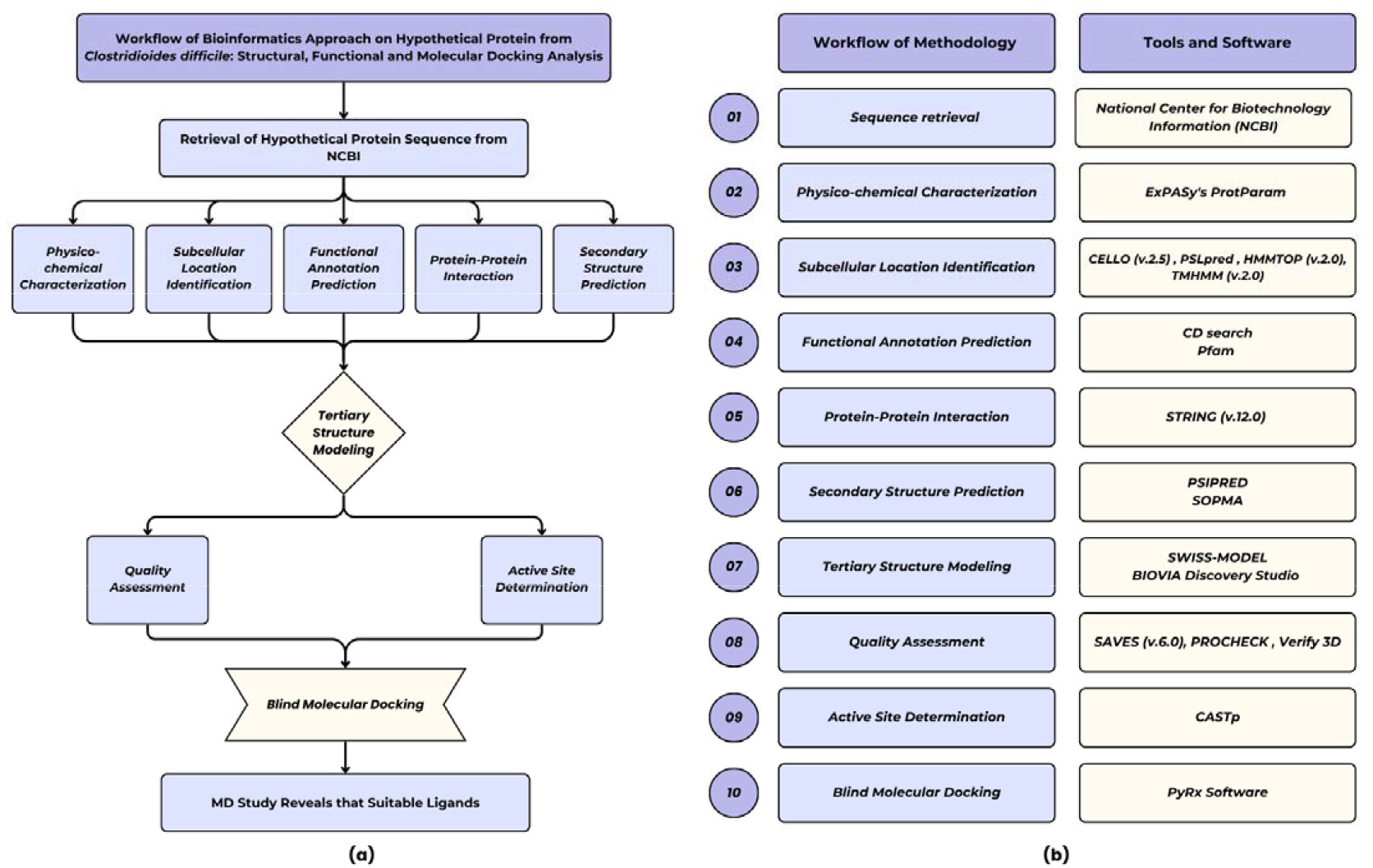
The complete workflow of methodolog **(a)** and bioinformatics tools **(b)** used in this study of Bioinformatics Approach on Hypothetical Protein from *Clostridioides difficile*.

### 2.1. Sequence retrieval

We retrieved the amino acid sequence of the hypothetical protein of *C. difficile* in FASTA format from the NCBI database using the accession ID WP_021373494 [11]. 1 The protein contains 65 amino acid residues.

### 2.2. Physicochemical characterization

We used the ProtParam tool of ExPASy to analyze the physical and chemical properties of the target protein, including molecular weight, aliphatic index (AI), extinction coefficients, GRAVY, and isoelectric point (pI) [12].

### 2.3. Subcellular location identification

We used the CELLO v.2.5 [13], PSORTb v.3.0.2 [14], SOSUI assessment tool [15], PSLpred server [16], HMMTOP v.2.0 [17], and TMHMM server v.2.0 [18] to document the subcellular location of the protein.

### 2.4. Functional annotation

We used the NCBI platform’s CD search tool to predict the conserved domain in the protein WP_021373494.1 [19]. We used the Pfam tool to determine the protein motif associated with the evolutionary relationships of the protein WP_021373494.1 [20].

### 2.5. Protein-protein interaction

We used the STRING v.12.0 program to determine potential protein-protein interactions (PPI) [21].

### 2.6. Secondary structure prediction

The secondary structure of WP_021373494.1 was predicted by the PSI-blast-based secondary structure prediction (PSIPRED) [22], and the self-optimized prediction method with alignment (SOPMA) framework [23] was used for element prediction.

### 2.7. Tertiary structure modeling

Currently, there is no experimentally concluded tertiary structure available for WP_021373494.1 of *C. difficile* in the Protein Data Bank (PDB). The 3D structure of the target protein was determined using the SWISS-MODEL server based on homology modeling [24]. The server automatically performs BLASTp search to identify templates for each protein sequence. The BIOVIA Discovery Studio Visualizer visualized the 3D model structure [25].

### 2.8. Quality assessment

We used the feature of the SAVES v.6.0 program to predict the Ramachandran plot and validate the predicted tertiary structure [26].

### 2.9. Active site determination

We used the Computed Atlas of Surface Topography of Proteins (CASTp) server to predict the active sites of the modeled protein [27].

### 2.10. Blind Molecular Docking

The AutoDock Vina program, utilized within PyRx, was employed to perform structure-based virtual screening by molecular docking. AutoDock Vina is a molecular docking and virtual screening tool that ensures accuracy in binding mode predictions by multithreading on multi-core platforms. It enables the analysis and prediction of ligand interactions with macromolecules. The ligands employed for docking were Phenylalanine arginine-β-naphthylamide (PAβN) and Carbonyl cyanide m-chlorophenyl hydrazone (CCCP) [28, 29]. The control inhibitors were obtained from the PubChem database. The docking outcomes were examined using Discovery Studio Visualizer.

## 3. Results

### 3.1. Sequence retrieval

We gathered the amino acid (AA) sequence of the hypothetical protein (WP_021373494.1) of C. difficile from the NCBI database. The protein contains 65 amino acids. Table **1** provides further information on the protein (WP_021373494.1).

**Table 1.**
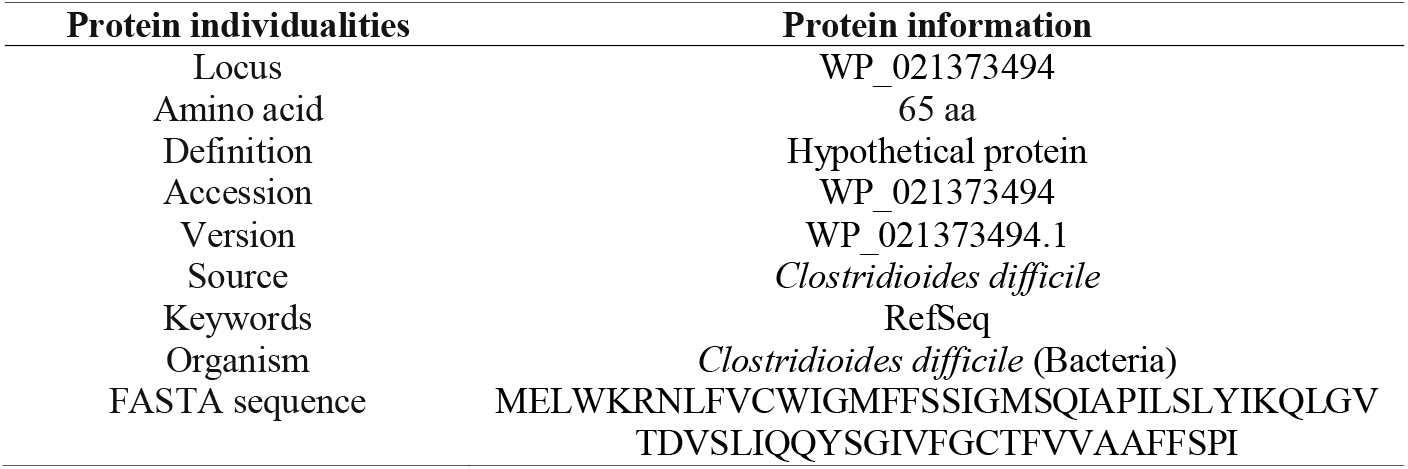
Hypothetical protein retrieval of *C. difficile* from NCBI.

### 3.2. Physicochemical characterization

The hypothetical protein (WP_021373494.1) contains a 65-amino-acid long sequence with a molecular weight of 7313.77 Dalton. The extinction coefficient (all pairs of Cys residues form cystines) is 14105. The protein is partially basic (pI 7.79), containing a total number of 2 negatively charged residues (Asp + Glu), and the total number of positively charged residues (Arg + Lys) is 3. The instability index, aliphatic index, and GRAVY are 56.55, 115.38, and 1.020, respectively, shown in Table **2**.

**Table 2.**
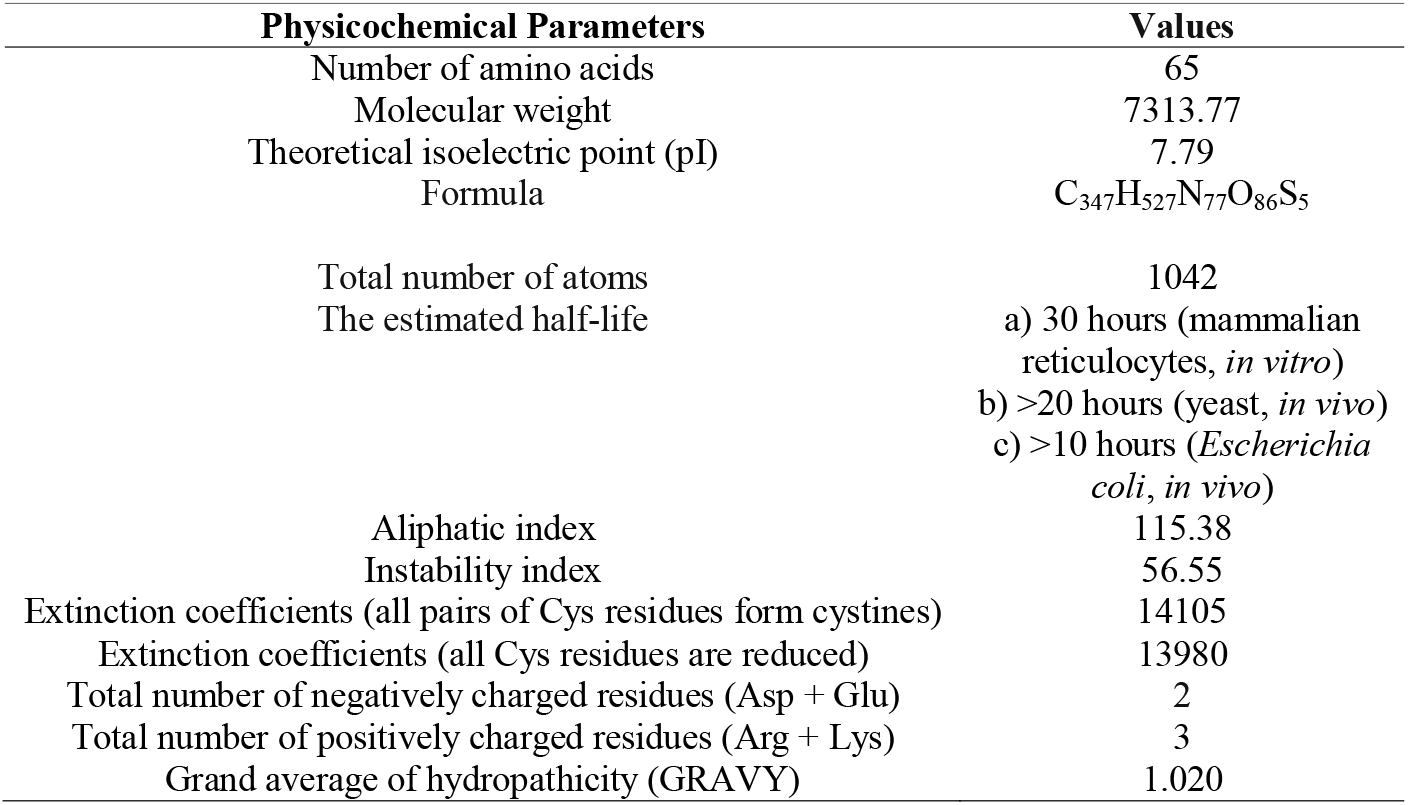
Physicochemical characterization of hypothetical protein of *C. difficile*.

| Physicochemical Parameters | Values |
| --- | --- |
| Number of amino acids | 65 |
| Molecular weight | 7313.77 |
| Theoretical isoelectric point (pI) | 7.79 |
| Formula | C<sub>347</sub>H<sub>527</sub>N<sub>77</sub>O<sub>86</sub>S<sub>5</sub> |
| Total number of atoms | 1042 |
| The estimated half-life | a) 30 hours (mammalian<br>reticulocytes, *in vitro*)<br>b) >20 hours (yeast, *in vivo*)<br>c) >10 hours (*Escherichia coli*, *in vivo*) |
| Aliphatic index | 115.38 |
| Instability index | 56.55 |
| Extinction coefficients (all pairs of Cys residues form cystines) | 14105 |
| Extinction coefficients (all Cys residues are reduced) | 13980 |
| Total number of negatively charged residues (Asp + Glu) | 2 |
| Total number of positively charged residues (Arg + Lys) | 3 |
| Grand average of hydropathicity (GRAVY) | 1.020 |

### 3.3. Subcellular location

We used the CELLO (v.2.5), PSORTb (v.3.0.2), SOSUIGramN, and PSLpred tools to assess the subcellular location of the protein (WP_021373494.1). The tools predicted the protein’s subcellular location as an inner membrane and cytoplasmic protein. The HMMTOP (v.2.0) and TMHMM (v.2.0) programs predicted that there were two transmembrane helices in the protein (WP_021373494.1) and emphasized the protein’s inner membrane and cytoplasmic location in the *C. difficle* shown in Table **3**.

**Table 3.** Subcellular localization of the hypothetical protein of *C. difficile*.

| Analysis | Result |
| --- | --- |
| CELLO (v.2.5) | Inner Membrane (1.662) and Cytoplasmic (1.426) |
| PSORTb (v.3.0.2) | Cytoplasmic |
| SOSUIGramN | Inner Membrane |
| PSLpred | Inner Membrane |
| HMMTOP (v.2.0) | 2 transmembrane helices |
| TMHMM (v.2.0) | 2 transmembrane helices |

### 3.4. Functional annotation

A protein domain is a conserved portion of a protein sequence that has a specific function and exists independently of the rest of the protein chain. The CD Search tool predicted a domain (accession ID: cl28910).This anticipated domain is associated with the Major Facilitator Superfamily (MFS), and Figure **2** shows the motif search tool Pfam and its predicted homologous superfamily, the Major Facilitator Superfamily (MFS) (accession ID IPR050497). This is a large and diverse collection of secondary transporters, including three uniporters, symporters, and antiporters.MFS proteins facilitate the transit of many compounds across cytoplasmic or inner membranes. These include sugar phosphates, medicines, neurotransmitters, nucleosides, amino acids, and peptides. They achieve this by using the electrochemical potential of the substrates being provided. Uniporters carry a single substrate, whereas symporters and antiporters move two substrates in opposite directions across membranes.MFS proteins typically include 400 to 600 amino acids and 12 transmembrane alpha helices (TMs) connected by hydrophilic loops. These proteins’ 11N- and C-terminal regions show minimal resemblance, which might be the result of gene duplication or fusion. Based on knowledge integration.We hypothesize that MFS proteins use a rocker-switch mechanism to alternatively access a single substrate. This is based on three kinetic investigations and the designs of various bacterial superfamily members, including GlpT (glycerol-3-phosphate transporter), LacY (lactose permease), and EmrD (multidrug transporter).Bacterial elements largely serve as nutrition absorption and drug efflux pumps, resulting in antibiotic resistance. Regarding numerous MFS proteins of medical relevance, such as the glucose transporter Glut4, which is impaired in type II diabetes, and the glucose-6-phosphate transporter (G6PT), whose mutation causes a glycogen storage disease.

**Figure 2.**
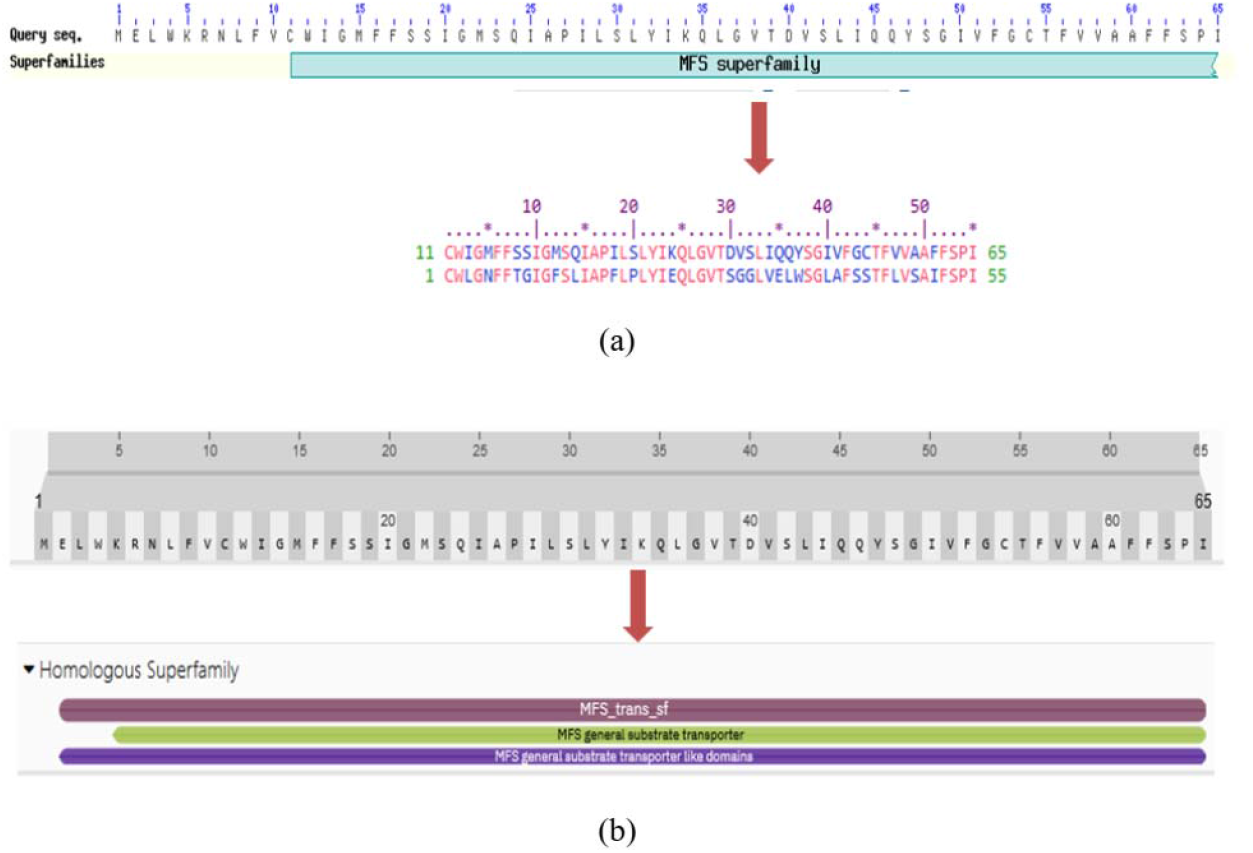
The hypothetical protein of *C. difficile* has undergone functional annotation. The graphical summary depicts the conserved domains detected in the query sequence. The aligned sequences indicate the conserved domains found on the query sequence when compared to the conserved protein domain family, MFS (CDD accession no.cl2890) (**a**). The Motif search and Pfam tools predicted the MFS homologous superfamily domain (accession no. IPR050497) of the protein WP_021373494.1 (**b**).

### 3.5. Protein-protein interaction analysis

The primary focus of protein-protein interactions is to acknowledge how cellular systems work. We used the STRING v.12.0 program to identify the protein-protein interaction (PPI) that occurred when MFS (CAJ67514.1) interacted with the 10 proteins shown in Figure **3**. The STRING program determined the functional protein companions with the interaction scores mentioned in Table **4**.

**Table 4.** List of proteins that interact with MFS (CAJ67514.1) identified through STRING.

| Interacting Proteins | Short Descriptions | Score |
| --- | --- | --- |
| cobA | Bifunctional uroporphyrinogen-III methyltransferase/uroporphyrinogen-III synthase. | 0.753 |
| CAJ67513.1 | Snoal-like polyketide cyclase. | 0.731 |
| CAJ67515.1 | Putative sodium:solute symporter; Belongs to the sodium:solute symporter (SSF) (TC 2.A.21) family. | 0.728 |
| CD630_06790 | Fragment of conserved hypothetical protein (C-terminal region). | 0.704 |
| CAJ67516.1 | Transcriptional regulator, LacI family. | 0.584 |
| CAJ67786.1 | Putative phage major capsid protein. | 0.537 |
| FprAnorR | Putative oxidoreductase; Experimentally verified as part of mature spore proteome | 0.533 |
| CAJ69808.1 | Putative phage major capsid protein | 0.509 |
| hpdB | Putative 4-hydroxyphenylacetate decarboxylase,catalytic subunit HpdB | 0.478 |
| ortB | 2-amino-4-ketopentanoate thiolase beta subunit | 0.456 |

**Figure 3.**
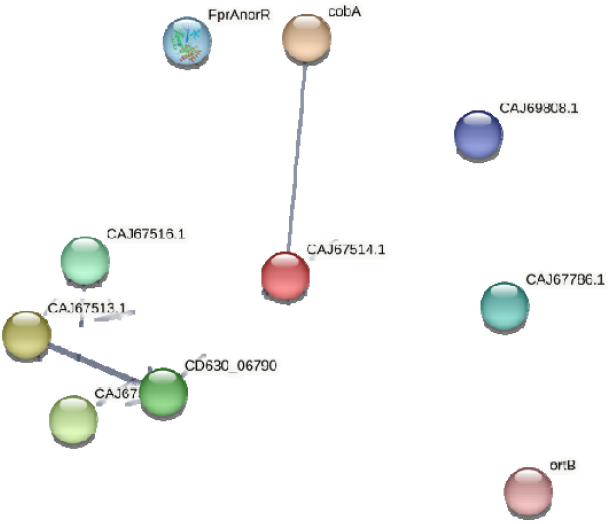
The STRING network of a protein regulates its interactions with other proteins. The first shell of interactors. For node content, empty nodes represent proteins with unknown 3D structures, whereas full nodes represent known or predicted 3D structures.

### 3.6. Secondary structure prediction

The PSIPRED software demonstrates increased confidence in predicting helices, strands, and coils (Figure **4a**). Table **5** presents the amino acid composition derived from the ExPASy ProtParam program. SOPMA utilized the default settings for simulating the secondary structure (Figure **4b**). It was forecasted that 61.54 percent of residues would be alpha-helices, but SOPMA estimated that 38.46 percent of residues would be random coils.

**Table 5.**
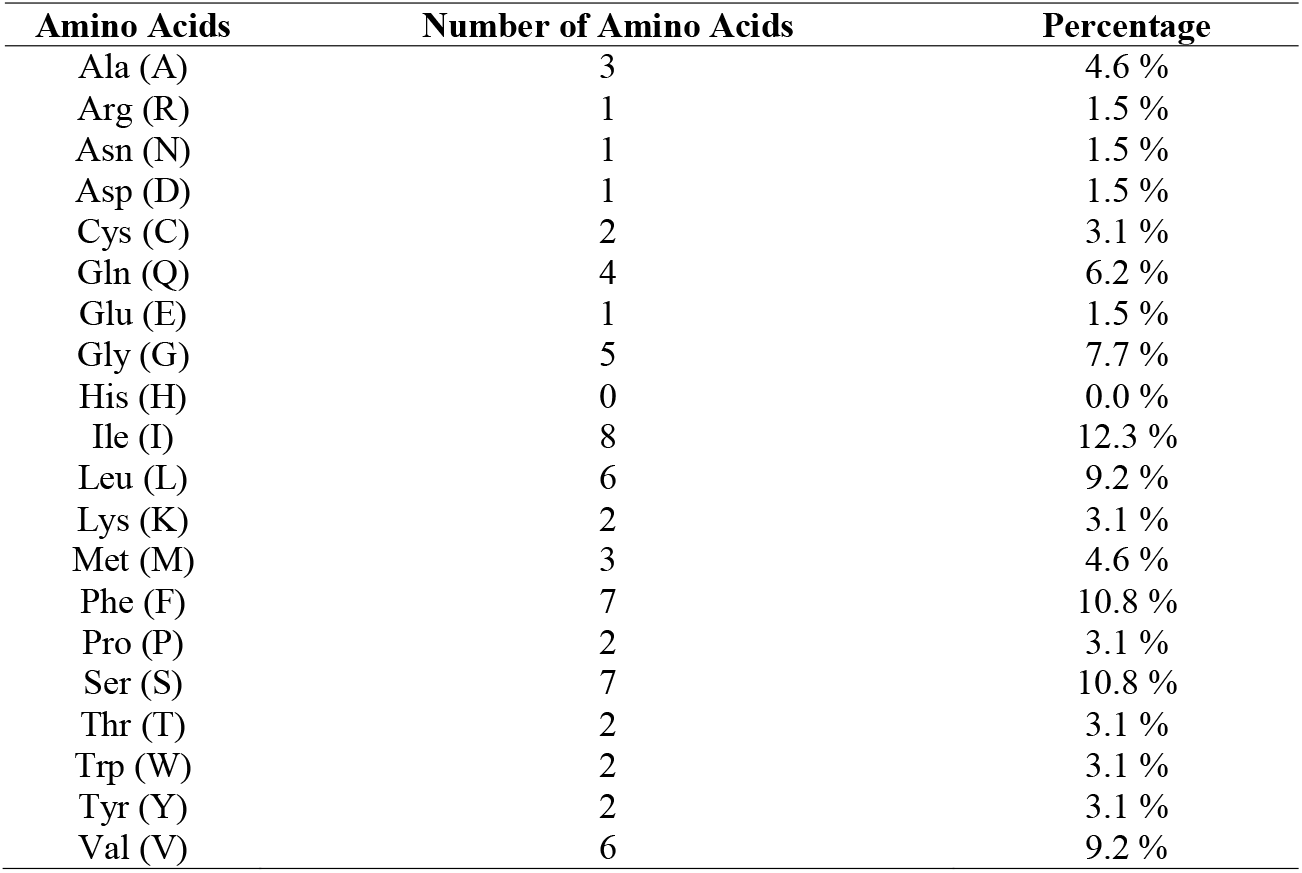
Amino acid composition of the hypothetical protein of *C. difficile*.

**Table 6.** Ramachandran plot statistics.

| Ramachandran plot statistics | Values (%) |
| --- | --- |
| Residues in most favored regions | 56 (100.0%) |
| Residues in additional allowed regions | 0 (0.00%) |
| Residues in generously allowed regions | 0 (0.00%) |
| Residues in disallowed regions | 0 (0.00%) |
| Number of non-glycine and non-proline residues | 56 (100.0%) |
| Number of end-residues (excl. Gly and Pro) | 2 |
| Number of glycine residues (shown as triangles) | 5 |
| Number of proline residues | 2 |
| Total number of residues | 65 |

**Figure 4.**
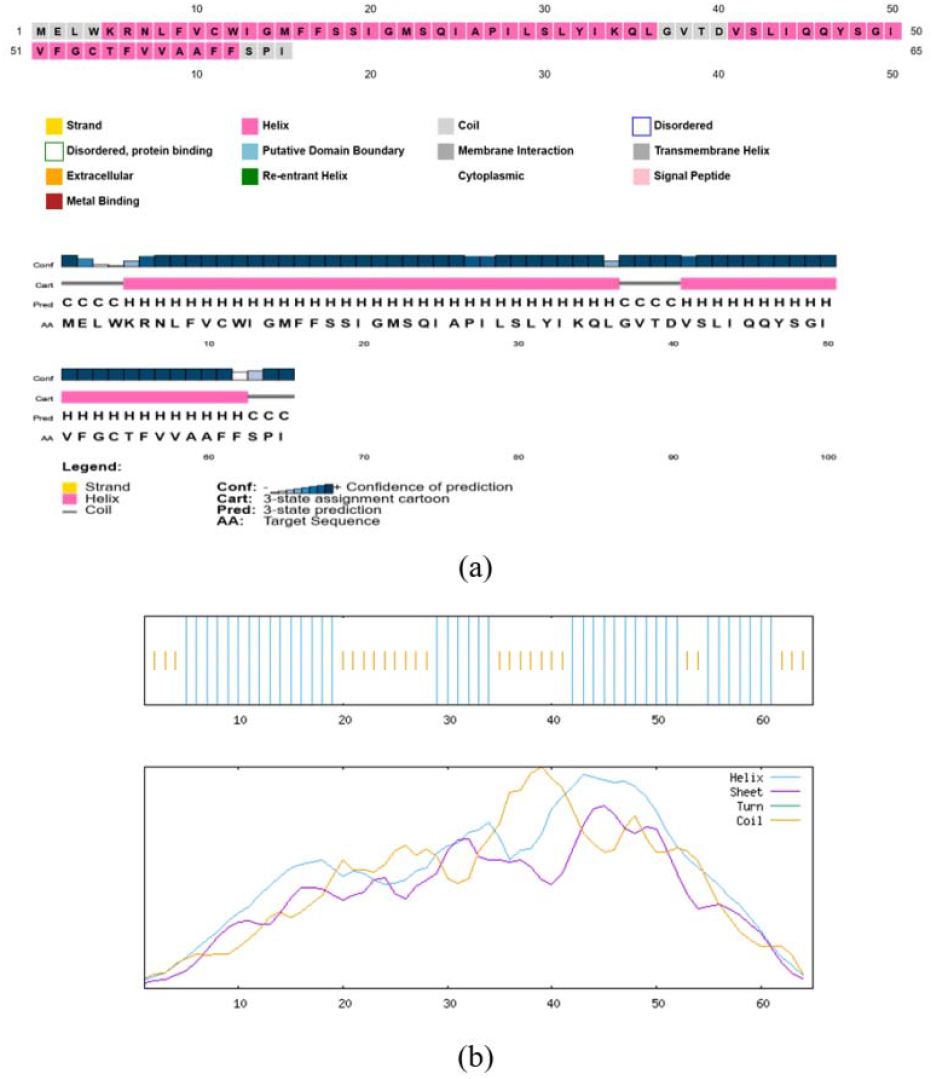
Secondary structure prediction of hypothetical protein of *C. difficile* by PSIPRED server (**a**) and SOPMA server (**b**).

### 3.7. Tertiary structure modeling

We obtained the tertiary structure of the protein from the SWISS-MODEL server using the template A0A125V656.1.A, demonstrating 95.38% sequence identity with the target protein. The modeling structure obtained through SWISS-MODEL is visualized in BIOVIA Discovery Studio software, which is depicted in Figure **5**.

**Figure 5.**
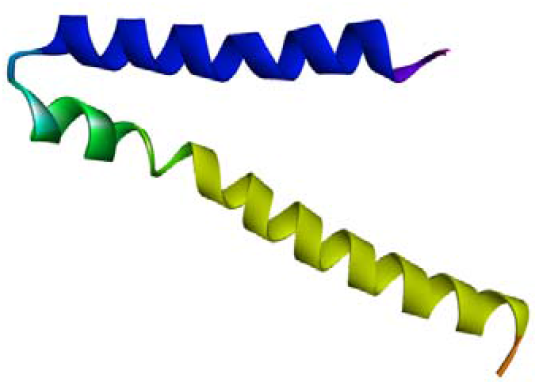
Predicted 3D structure of the target protein through the SWISS-MODEL server visualized in BIOVIA Discovery Studio software.

### 3.8. Quality assessment

We used the PROCHECK program on the SAVES server to check the structure quality of the modeled protein. The ψ angle and the φ angle are exhibit in Figure **6**. Residues in the most preferred regions engulfed a total of 100.0%, which validated the protein’s modeled tertiary structure (WP_021373494.1). Also, there were no residues in the supplementary allowed regions, generously allowed regions, or disallowed regions (0.00%). There were 56 non-glycine and non-proline residues, as well as 56 end residues (excluding Gly and Pro), which is 100%. The protein 3D structure mentioned in Table **7** contained 5 residues of glycine and 2 residues of proline.

**Table 7.**
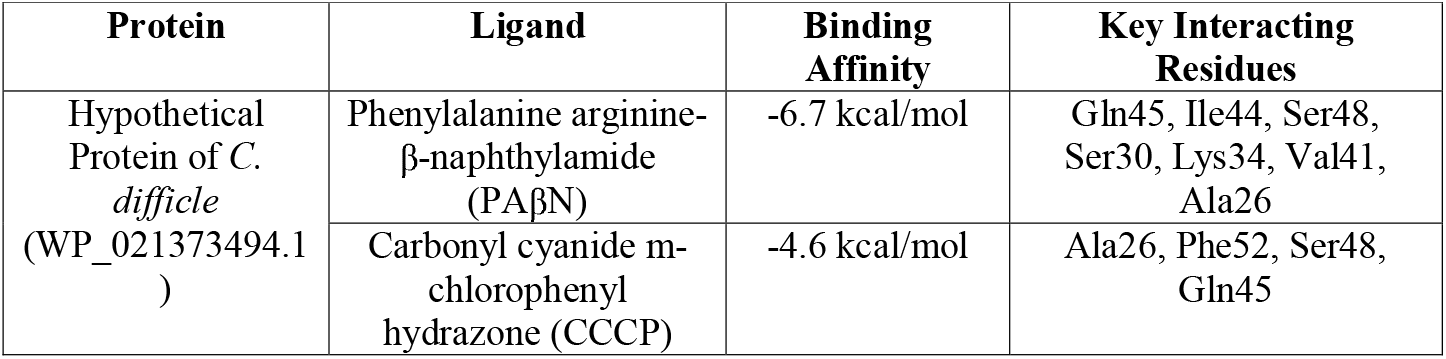
Summary of docking analysis results from Autodock Vina tool, involved in PyRx.

**Figure 6.**
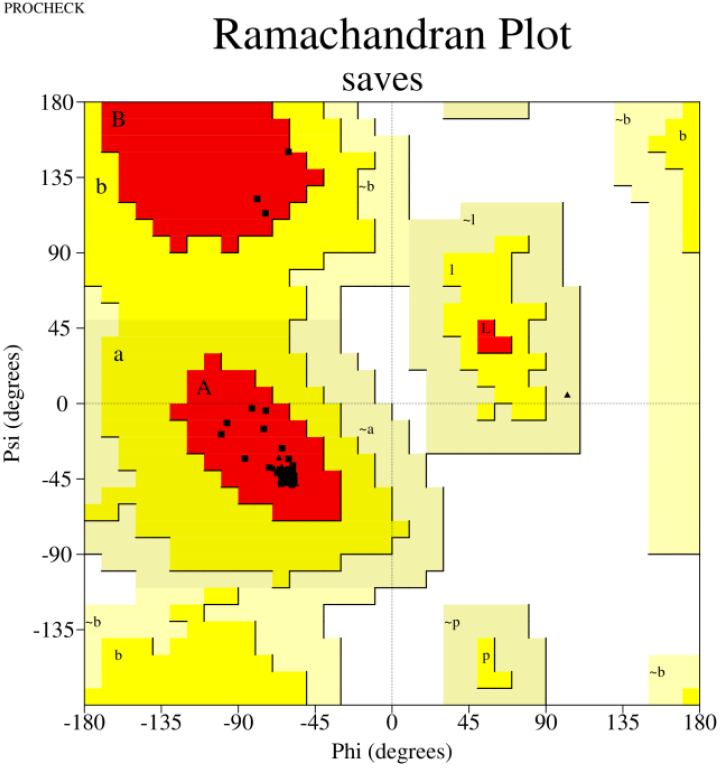
The PROCHECK tool confirmed the Ramachandran plot statistics for the three-dimensional structure that was modeled.

### 3.9. Active site determination

The program, known as the Computed Atlas of Surface Topography of Proteins (CASTp) v.3.0, predicted five distinct active sites for the modeled protein. Therefore, this study employed the CASTp server to analyze the protein’s active site. Figure **7** illustrates the region involved in active site formation.

**Figure 7.**
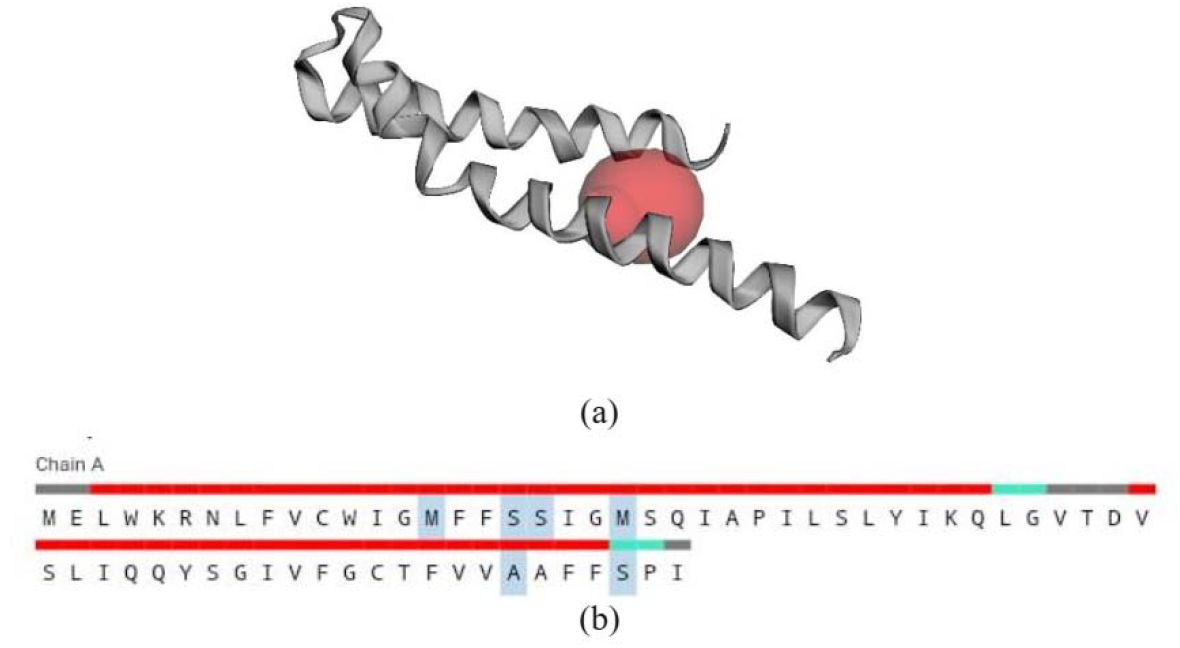
The “red sphere” represents the protein’s active sites **(a)** and also the blue color of amino acid regions represents the active site of the hypothetical protein of *C. difficile* (**b**).

### 3.10. Molecular Docking Analysis

A docking study of the target protein and the ligand was conducted with the AutoDock Vina program included in the PyRx software. The ligands phenylalanine arginine-β-naphthylamide (PAβN) and carbonyl cyanide m-chlorophenyl hydrazone (CCCP) were docked with the target putative protein of C. difficile (WP_021373494.1). A robust binding affinity was noted for the ligand with both proteins. The binding affinities of phenylalanine arginine-β-naphthylamide (PAβN) and carbonyl cyanide m-chlorophenyl hydrazone (CCCP) for the model protein were -6.7 kcal/mol and -4.6 kcal/mol, respectively, with a shared grid box center at X: 0.4849; Y: -0.1480; Z: -1.1136 and dimensions (angstrom) of X: 43.2553; Y: 30.8400; Z: 48.3064. The docking analysis reveals that the phenylalanine arginine-β-naphthylamide (PAβN) ligand binds with specific amino acids like Gln45, Ile44, Ser48, Ser30, Lys34, Val41, and Ala26 with -6.7 kcal/mol binding energy, while the carbonyl cyanide m-chlorophenyl hydrazone (CCCP) ligand binds with specific amino acid residues like Ala26, Phe52, Ser48, and Gln45 with -4.6 kcal/mol binding energy. Strong hydrogen bonds and the docked complex’s lowest energy value indicate a solid link between the ligand and receptor molecule.The findings of the docking analysis are showed in Table **7** and Figure **8**.

**Figure 8.**
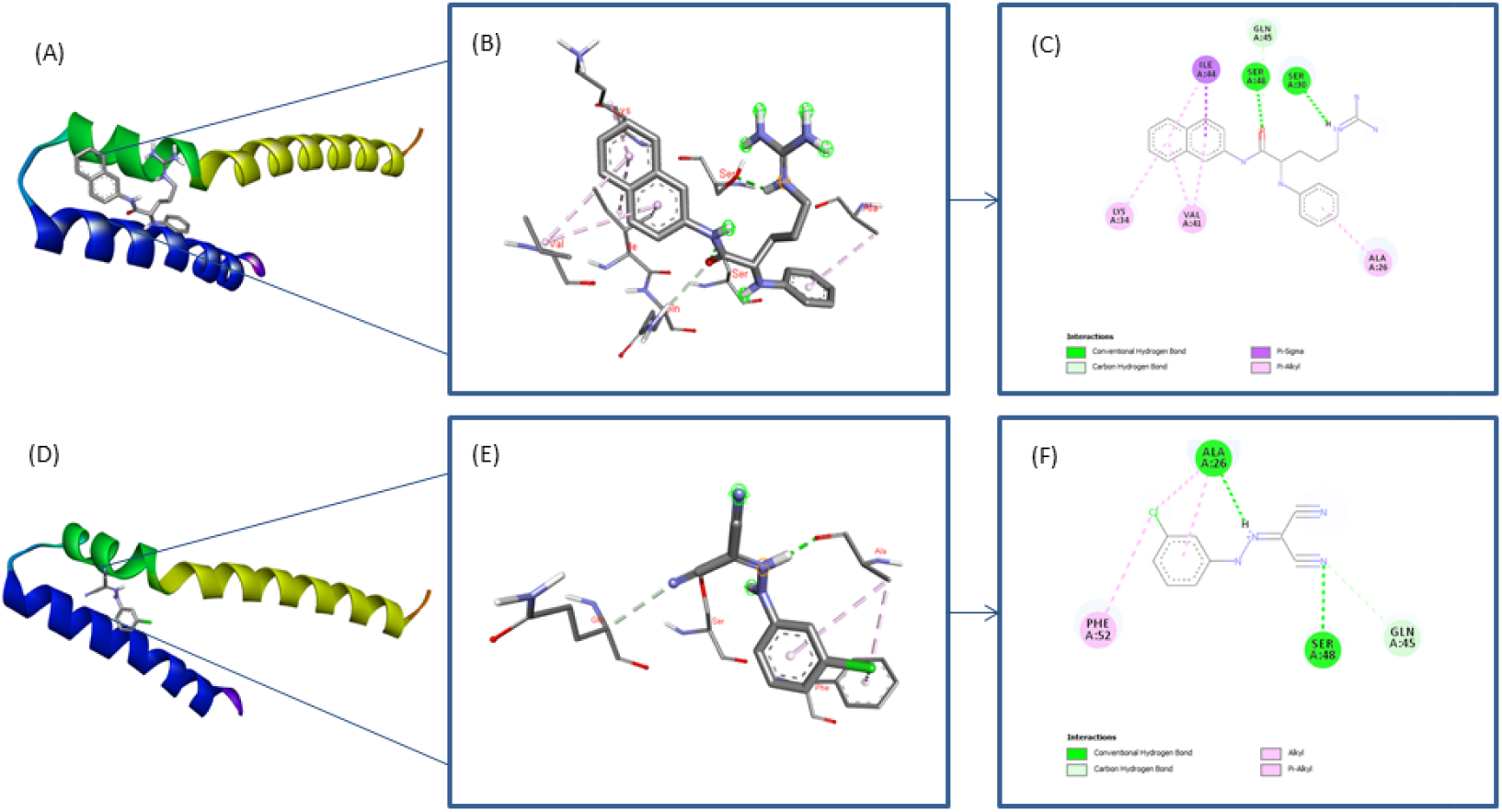
Phenylalanine arginine-β-naphthylamide (PAβN) and carbonyl cyanide m-chlorophenyl hydrazone (CCCP) with hypothetical protein of *C. difficle* (WP_021373494.1): phenylalanine arginine-β-naphthylamide (PAβN) bound with protein in 3D structure **(A)**, ligand (PAβN) interplay with the protein (WP_021373494.1) **(B)**, key interacting residues of the protein (WP_021373494.1) with ligand (PAβN) in 2D structure **(C)**, and carbonyl cyanide m-chlorophenyl hydrazone (CCCP) bound with protein in 3D structure **(D)**, ligand (CCCP) interaction with the protein (WP_021373494.1) **(E)**, key interacting residues of the protein (WP_021373494.1) with ligand (CCCP) in 2D structure **(F)**; all docking results analyzed by BIOVIA Discovery Studio Visualizer.

## 4. Discussion

The hypothetical protein (WP_021373494.1) of *C. difficile* obtained from the NCBI contained 65 amino acids in its sequence. The physicochemical values determined the protein’s partial basic nature (pI 7.79). Besides solubility, the pI can influence molecular behavior in chromatography and electrophoresis [30, 31]. These strategies depend on molecules being separated by their charge and size, and the pI can be used to guess how molecules will move around in these systems [30, 32]. Moreover, the positive GRAVY (1.020) indicated the protein was hydrophobic. Three molecules of Arg and Lys in the sequence may influence the GRAVY value, as the quantity of positively charged AA was higher than that of negatively charged (Asp and Glu).

The subcellular localization (SL) of a protein is essential for its function since it dictates the protein’s capacity to engage with particular molecular partners and partake in specific biological processes [33, 34]. Proteins are frequently confined to particular cellular compartments where their functionality is necessary [33, 35]. The protein was predicted to be involved in the inner membrane and cytoplasm via the SL investigation. The inner membrane (IM) and cytoplasmic proteins (CPs) have unique yet interrelated functions in cellular activity. IM proteins, situated within membranes such as the mitochondrial IM or the plasma membrane, are crucial for functions including transport, signal transduction, and energy production [36, 37]. They frequently function as channels, pumps, or receptors, governing the exchange of chemicals and signals across the membrane. CPs function within the intracellular fluid, facilitating enzymatic processes, cytoskeletal structure, and molecular signaling [38, 39]. These proteins facilitate critical functions like metabolism, cellular division, and stress responses. The functional interaction between the IM and CPs is essential for sustaining cellular homeostasis, as several CPs engage with or control IM components to synchronize cellular responses to environmental and physiological alterations [33,40]. Comprehending the functions of these proteins elucidates essential biological processes and their ramifications for health and illness [41, 42].

Subsequent functional analyses revealed that the protein was part of the MFS group, implicating its role in function via a rocker-switch mechanism [43]. The PPI interaction network identified its link with an additional 10 proteins that interacted with this MFS protein. The secondary structural level results from hydrogen bonding among the backbone atoms of amino acids and acts as a fundamental component of the protein’s overall three-dimensional structure [44, 45]. Secondary structures confer both stiffness and flexibility in designated locations, facilitating proteins in executing their functions proficiently, such as establishing active sites in enzymes or binding surfaces in receptors [46, 47]. Disruptions in secondary structure may result in protein misfolding and aggregation, potentially contributing to illnesses. Moreover, examining secondary structures improves our comprehension of protein design, folding mechanisms, and interactions, which are essential for drug discovery and the creation of biomimetic materials [48, 49]. Consequently, the secondary structure is essential for clarifying the complex link between a protein’s structure and function [49, 50]. The analyses revealed that 61.54% of the protein participated in the alpha-helix formation, which was almost 23.08% higher than the value of the random coil (38.46%).

The tertiary structure (3D) of a protein is essential for its biological activity, as it denotes the 3D configuration of AA residues that enables the protein to achieve a functional shape [51]. This structure is reinforced by several interactions, including hydrogen (H) bonds, hydrophobic contacts, ionic bonds, and disulfide bridges (DBs), which collectively dictate the protein’s conformation and dynamics. The right way to fold the 3D creates active sites, binding pockets (BPs), and interaction surfaces [52,53]. These are necessary for enzyme activity, chemical recognition, and signal transduction (ST). Disruptions or misfolding of 3D can compromise protein function, resulting in illnesses [53, 54]. Comprehending 3D is essential for elucidating protein functions, building functional biomolecules, and developing medications that target specific structural motifs, thus improving treatment techniques and advancing biotechnological applications [53, 55]. The protein’s 3D structure was constructed utilizing the most appropriate template, exhibiting 95.38% similarity. Structural quality assessment documented that all sequences (100%) were in the most favored regions. According to Ramachandran plot analyses, there was no sequence in the additional, generously, or disallowed regions.

Molecular docking (MD) analysis is a computer method employed to forecast the interaction between molecules, usually a tiny ligand and a target protein, to comprehend their binding affinity and interaction manner [56]. This approach is essential in drug discovery and design (DDD), as it offers insights into the binding interactions between a possible drug candidate and its biological target, such as an enzyme or receptor [57, 58]. Through the simulation of the docking process, researchers may discern critical interactions, including hydrogen (H) bonds, hydrophobic contacts, and ionic interactions, that stabilize the ligand-protein (L-P) complex [59]. MD makes it easier to sort compounds by their binding scores, which helps choose the most promising ones for further testing in the lab. It diminishes the time and expenses linked to experimental screening and aids in optimizing medication creation by directing structural alterations to improve efficacy and specificity [60, 61]. MD analysis is an essential instrument in contemporary computational biology and pharmacology. The MD study was performed based on the protein’s active sites so that the selected ligand molecules could bind to the binding pocket with lower energy scores [62, 63]. PAβN interacted with the protein with minimal interacting energy (– 6.7 kcal/mol) compared to another ligand, CCCP (– 4.6 kcal/mol).

## 5. Conclusions

*C. difficile*, a pathogen linked to severe infectious colitis, is now present in Asia, resulting in considerable illnesses and fatalities globally. The recent rise in C. difficile prevalence, together with the emergence of more virulent strains in Europe and North America, certainly suggests a same trend in Asia. The study aims to look at many aspects of the protein, such as its physicochemical properties, where it is found within cells, how it works, how it interacts with other proteins, how its structure can be predicted and confirmed, and how active sites can be found for possible ligands. The protein exhibits partial basicity and hydrophobicity, as indicated by the examination of its physicochemical properties. The protein exhibits functions in the inner membrane (IM) and cytoplasm, characterized by two transmembrane helices. Furthermore, the protein acts as a secondary transporter within the MFS system, facilitating the translocation of various molecules across cytoplasmic or interior membranes. To uncover novel therapeutic medicines, we targeted the protein’s active areas as possible ligand-binding sites. The MD investigation examined the interaction between the selected ligands (PAβN and CCCP) and the protein. PAβN was determined to be a more efficacious ligand than CCCP owing to its reduced energy demand for protein interaction. Although it’s potential as a therapeutic target, the study’s reliance on computational predictions underscores the need for and experimental validation. Further investigations are essential to confirm the protein’s biological role and therapeutic significance.

## Institutional Review Board Statement

Not applicable.

## Informed Consent Statement

Not applicable

## Data Availability Statement

All research data are provided in the current article.

## Acknowledgments

Each writer contributed equally to this work. Thank you to the coauthors for offering adequate assistance, financial support, editing, and writing to assure the study’s success. This research work was assisted by the NHMRC Investigator Grant (APP1175047) for A.A.I.S.

